# A large-scale crowd-sourced annotated acoustic dataset of Indian fauna

**DOI:** 10.64898/2026.07.20.739496

**Authors:** Vijay Ramesh, Sameer Singh, Paul Pop, Pooja Choksi, Prabhjeet Singh, Sarika Khanwilkar, Sonal Teotia, Pralhad Burli, Kadambari Devarajan, A Ashwin, Amisha A Nakhwa, Mohammad Abdus Shakur, Rohan Baishya, Nishant Bhagwat, Siddharth Biniwale, Chiranjib Bora, C S Swathi, Nibedita Chakraborty, Senan D’Souza, Elrika D’Souza, Ram Dayal Vaishnav, Kadambari Deshpande, Aavika Dhanda, G Abhishaek, Aarini Ghosh, Rajkamal Goswami, K N Abhinav, K P Nidhichand, Bittu K Rajaraman, K V Gururaja, Vandana Kannan, Vijay Karthick, Mayuri Kotian, Harsha Kumar, Punnen Kurian, Malavika Madhavan, Komal Meena, Ayesha Mohammad Maslehuddin, Pravar Mourya, Madhushri Mudke, R J Pradeeshwar, Rohit R S Jha, Kavya Ramesh, S Sangeeth Sailas, Taksh Sangwan, Siddharth Mahesh, Ryan Satish, Anusha Shankar, Cimila Sibichen, Ramit Singal, Gaurav Singh, Navneet Singh, Iqbal Singh Bhalla, Navaneeth Sini George, Meghana Srivathsa, Pavithra Sundar, Syeda Tabassum Tasfia, Alayna Trejo, Yana V Rasheva

## Abstract

Global rates of biodiversity loss warrant conservation action and monitoring at large geographic scales. Conservation technologies such as acoustic monitoring in conjunction with deep learning now enable us to monitor wildlife simultaneously across space and time. However, for a significant proportion of biodiversity in tropical regions, we cannot yet rely on automated recognition approaches because we lack acoustic templates to robustly train deep learning algorithms. In this paper, we relied on a novel participatory approach, enlisting researchers, conservation practitioners, and nature enthusiasts to create a unique crowd-sourced, open-access dataset of acoustic annotations across taxonomic groups for biodiversity in India. Our dataset comprises 3311 minutes of ‘strongly labelled data’ (bounding boxes or annotations for a species vocalization) and 2504 minutes of ‘weakly labelled data’ (indicating the presence of a species within an audio file but lacking bounding boxes) for 518 species across India, spanning 25 of 36 states and union territories. We present metadata and code for data processing and highlight the strengths of a participatory approach to biodiversity monitoring.

## Background & Summary

Biodiversity loss is occurring at unprecedented rates globally ^1,2^, and urgent action is required to implement effective wildlife monitoring approaches at scale to ensure robust conservation. The use of novel conservation technologies and advances in deep learning now allow conservation scientists and practitioners to monitor wildlife rapidly, potentially preventing and stemming further species declines^3^. Acoustic monitoring in particular, has gained significant traction for detecting species threats, studying species behavior and habitat use, and assessing the impacts of anthropogenic changes on biodiversity^4^. Currently, the pipeline for the use of acoustic monitoring for species-specific assessments predominantly involves the use of an acoustic recorder (passive or focal) at an ecological monitoring station or site, followed by analysis of audio to identify specific species. Historically, species-specific acoustic analysis involved manually listening to large amounts of acoustic data to identify the target species. As training data in the form of acoustic templates became available for a larger number of species, semi-automated classifiers and detectors were built to identify the species of interest ^5^. With the advent of artificial intelligence and its reliance for biodiversity monitoring, the field of bioacoustics is rapidly shifting towards using automated species recognition approaches that rely on deep learning models. Although widely used, these deep learning models (e.g. BirdNET, Perch etc.) are currently constrained by the quantity and quality of available training data, which significantly affects their performance ^6,7^.

Tropical regions, which support disproportionately higher levels of biodiversity than temperate latitudes^8^, are locations where existing deep learning models perform poorly for several reasons. This is primarily due to a lack of species annotations and the inability to distinguish species vocalizations from noisy backgrounds. While advances in deep learning, such as agile modeling, embedding search, transfer learning, and few-shot learning, help overcome existing limitations, we still require robust annotations of the target species to achieve high performance^9,10^. The creation of training and benchmark datasets for tropical biodiversity will enable conservation scientists and practitioners to train robust classifiers and evaluate their performance effectively. Until now, efforts to create such annotated datasets have been taxonomically and geographically biased. For example, BirdNET and Perch have been trained primarily on annotated acoustic data from temperate regions of the Global North^6,7^. Further, existing annotated datasets are predominantly available for birds and anurans^11^ (to a limited extent) from specific sites in North America, Europe, and parts of South America, leaving large gaps for highly biodiverse countries, such as India. In addition, there is a geographic imbalance in such annotated data, with a large proportion sourced from a single/few ecosystems and regions of the world, thereby reducing the model’s ability to learn from acoustic features across the wider geographic range of a species’ distribution^12^.

To create annotated data across taxonomic groups for tropical biodiversity, we need a scalable approach to data collection that can engage a broad audience interested in nature sounds. For several decades, community scientists have contributed significantly to the identification and distribution of biodiversity^13^. With the advent of public-engagement and participatory science platforms, such as eBird and iNaturalist, the number of people engaging with nature and contributing their observations to centralized platforms has increased exponentially^14,15^. Across India, researchers, conservation practitioners, sound recordists, government officials, wildlife enthusiasts and naturalists, and the general public banded together to co-create the India Ecoacoustics Network (IEN; https://indiaecoacousticsnetwork.weebly.com/) in 2024. This platform serves as a communal space for knowledge exchange and capacity building — uniting people through the shared goal of using sounds to monitor, protect, and better understand India’s rich biodiversity. Here, we provide the first crowd-sourced, open-access dataset of acoustic annotations across taxonomic groups for biodiversity in India. As part of this effort, members of the IEN contributed acoustic data for 518 species of birds, mammals, reptiles, amphibians, insects, freshwater, and marine taxonomic groups across India. This crowd-sourced, open-access effort enables species monitoring by providing acoustic templates for training deep learning models.

## Methods

### Informational surveys and minimum data requirements

In May 2025, an informal survey was administered to gauge interest from those within and beyond the IEN who possess acoustic data and are willing to contribute to an open-access dataset of acoustic annotations of biodiversity within India. This survey was circulated via the IEN WhatsApp group, the IEN and other email lists, and through personal connections. We received responses from 99 participants, who helped refine the aims and goals of the effort. Participants posed questions about data use, data privacy and sharing, and contributor effort and acknowledgements.

Following this initial informal survey, we created an FAQ document outlining the goals of our effort and answering questions about the collation of the open-access dataset. Additionally, this FAQ clarified questions regarding minimum data contribution for co-authorships (30 minutes of audio recordings), data privacy, management, publication, and long-term storage. We requested contributors to provide one or two types of data. Type A data refers to strong labels with bounding boxes associated with a target species within an audio file. Type B data refers to weak labels or a recording that contains a target species without any boxes drawn around its presence in that audio file.

We requested that all submitted data identify the taxonomic group at the species level. However, in rare cases where limited annotated data were available for certain taxonomic groups, such as insects, we accepted identifications at the highest possible rank. There was no specification on the minimum or maximum number of species that could be present within a single audio file. At the very least, we requested that each submitted audio file be 10 seconds in length, with a maximum of 20 minutes, for ease of processing. The FAQ was reviewed and iterated by a core team of eight co-authors and is provided as supplementary material.

### Data sharing agreements and data submission

We circulated a consent form to all contributors interested in participating in this effort, along with the FAQ document. We shared this form widely through the IEN communication channels, email lists, social media, and word of mouth and personal connections. The consent form also provided a link for the contributor to upload their data. The uploaded data could come from a variety of sources, including mobile phones, dedicated hand-held focal recording devices, and passive acoustic monitors. All audio data formats were acceptable for submission, but we recommended that the data be in one of the following formats: .wav, .flac, or .mp3.

In the data submission form, we provided specific instructions for submitting Type A and Type B data. For Type A data, we provided a template .xlsx file with three sheets. The first sheet contained metadata for the submitted media file, including the file name, the geographic location (the recording’s latitude and longitude, which could be hidden upon request by the authors), the recording date, and the recording type (active or passive). The second sheet contained columns for start and end times and lowest and highest frequencies of the annotations/bounding boxes identified in the audio file, along with the organism’s scientific name. The third sheet for Type A in the .xlsx file requested information on the authors, including their email IDs and affiliations. For Type B, we requested the same information as above, except for the second sheet pertaining to annotations.

Once authors had filled out the sheets in template.xlsx for Type A and Type B data, we asked contributors to provide a single zipped file for each data type containing the template.xlsx and all corresponding audio files. All the data submitted was initially stored in a folder for processing prior to curation for Zenodo. In addition to providing an easy-to-access link for submitting data, P.P., S.S., S.K., and S.T. created detailed tutorials for annotation of Type A data using Raven Pro 1.6.1^16^, automation scripts for data entry for both Type A and Type B data, and ran virtual and in-person workshops for interested participants between September 2025 and November 2025. The annotation protocol, consent form, and data submission form are provided as supplementary material.

### Data processing

We processed the submitted data (for both Type A and Type B) in three distinct stages: parsing, standardization, and transformation using an automated data processing pipeline (accessible via the GitHub repositories associated with this paper)(Figure 1). Although the expected metadata format and file structure were outlined in the form, the majority of submitted zipped folders included template.xlsx files that required manual edits and fixes before further data processing.

**Figure 1.**
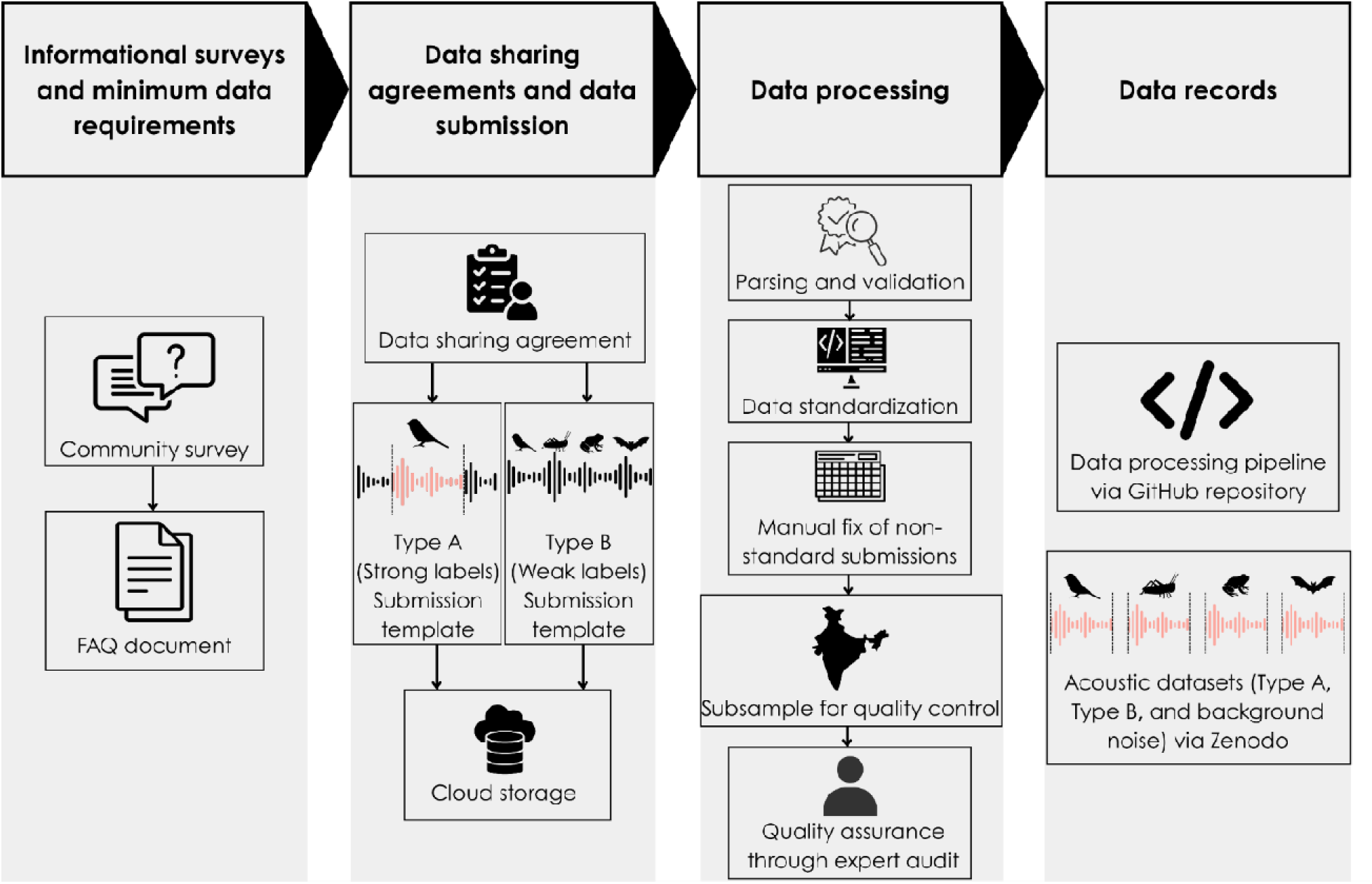
An overview of the methodological workflow for the creation of a crowd-sourced, open-access acoustic dataset of Indian fauna.

In the parsing step, we verified the presence of mandatory columns for all rows across all metadata sheets in the submitted .xlsx files (for Type A and Type B data, respectively). After parsing, if all mandatory columns were present, we unzipped and copied the audio files, the metadata sheet, the author information sheet, and the annotation sheet (only for Type A) into dedicated cloud storage destination folders for further processing. We renamed all audio files using binary hashing, an algorithmic procedure that replaces each filename with a unique alphanumeric string to avoid duplicate filenames across submissions. At this step, we also began maintaining a primary tracking table in a central database server for each data type to support downstream processing.

Once we parsed and saved all the submitted data in an unarchived format in the cloud, we standardized and augmented each record for further analysis. This step included standardizing scientific names and attaching their relevant usageKey using the Global Biodiversity Information Facility (GBIF) backbone taxonomy^17^. We utilized the pygbif Python interface (v0.6.5) to programmatically query the GBIF API (queried in January 2026). For names identified as synonyms, we followed the acceptedUsageKey to resolve them to their currently accepted scientific names as recognized by the GBIF backbone. We also matched each record’s latitude and longitude to an Indian state or union territory using an administrative boundary layer.

The submitted .xlsx files contained various typographical errors across multiple columns, including the scientific name. In the data transformation stage, we used fuzzy logic to identify and group all misspelled data pertaining to the identified organism into a single header, and to collate all environment- and vocalisation-related information into a single field named notes in the final published dataset. At the end of this stage, we had two metadata schemas (one for Type A and one for Type B), an annotation schema for Type A, and a consolidated authors schema for all contributors. Please note that our initial data processing missed a few errors, which were subsequently fixed during technical validation (see below).

We created primary keys in both Type A and Type B schemas, ensuring that the same media was not double counted across submissions. For the Type A metadata schema, we used the media file’s binary hash as the primary key. However, for Type B, we used a combination of binary hash and the scientific name to maintain a separate record for each identified organism and file. The records in the annotations schema were made unique by using a combination of scientific_name, begin_time, end_time, high_freq, and low_freq as the primary key. We used the authors’ email addresses to make author records unique. All these schemas were hosted on a PostgreSQL (v16.11) database running on an Amazon Web Services virtual machine and were served via Mathesar (v0.8.0; https://mathesar.org/), which provided a collaborative interface for the co-authors to review and analyse before exporting it to Zenodo.

### Data Record

The audio data and associated metadata are available under the Creative Commons license (CC-BY-NC) and are deposited in Zenodo. After parsing, verifying, and transforming the data into a standard format, the final dataset created contained contributions from 58 authors for 518 species (11 unique species identified only to genus, family, class, or order level), spanning 25 out of the 36 states and union territories of India (Figure 2). Type A data originated from 22 of the 36 states and union territories of India, whereas Type B data originated from 19 of the 36. The published dataset, as of March 2026, contains 3311 minutes of Type A and 2504 minutes of Type B data.

**Figure 2:**
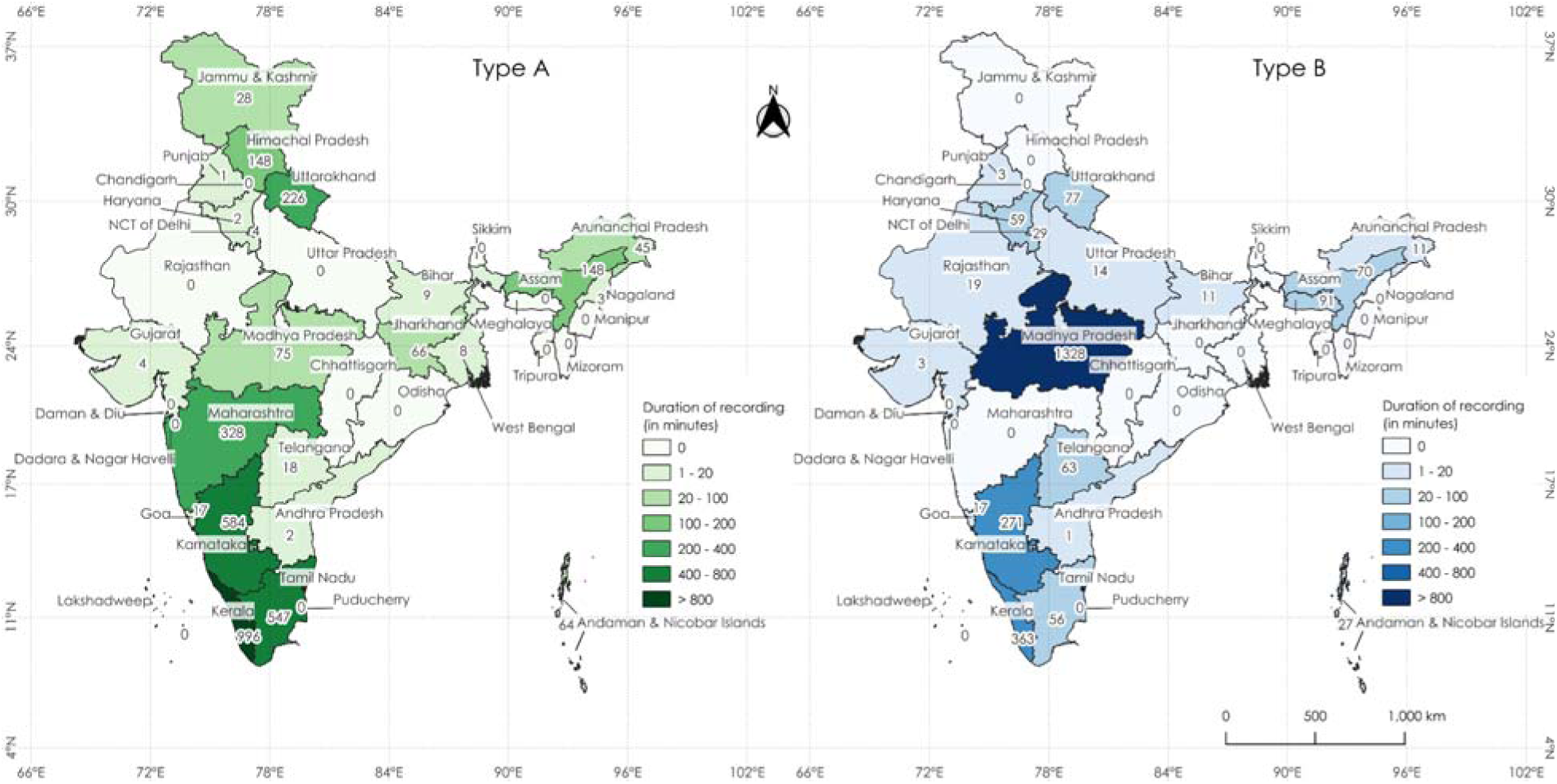
Chloropleth maps showing the duration of recordings (in minutes) across India, for both Type A and Type B data. Two Type B recordings (around 71 seconds in total media length) with no geographic information are not included in the display. Map not for scale.

Our dataset is organized as follows on Zenodo: (i) Type A data (with a sub-folder for audio, a metadata.csv file, and an annotations.csv file); (ii) Type B data (with a sub-folder for audio and a metadata.csv file); (iii) Background (with a sub-folder for audio, metadata.csv, and an annotations.csv file). Data upload size limitations on Zenodo prevent us from making a single record. Hence, the overall dataset is split in Zenodo into three records corresponding to the above data organization. For ease of user access, the audio folder (for both Type A and Type B) was further divided into subfolders, with audio files organized by state (geographic designation).

Please refer to the Zenodo description for more details.

The following list describes information contained within the columns of metadata.csv file for Type A data, along with any data naming conventions that may be followed:

● media_file_name: the unique file name corresponding to the audio file. We generated the following unique identifier for each file: **{sequential_fileID}_{state_name}_{recording_date}_{alphanumeric_string}.wav.** For example, 0001_arunachalPradesh_20200201_4e432694.wav. We relied on this naming convention to ensure no duplicates were generated and to ease data organization for the Zenodo upload.
● recording_date: the date of recording in YYYY-MM-DD format.
● longitude: data presented in decimal degrees. If the data was hidden or unavailable, the column contained an NA.
● latitude: data presented in decimal degrees. If the data was hidden or unavailable, the column contained an NA.
● author_names: names of all contributing authors for this media file separated by a semi-colon.
● email_addresses: all contributing author email addresses separated by a semi-colon.
● notes: supplementary information provided by the authors. This field typically contains environment, weather, and habitat information when present in the metadata.csv.
● archive_name: the name of the zipped folder containing the audio files. Since we organized audio by geographic state, this column referenced the state name. For example, typeA_audio_himachal_pradesh.zip would contain all audio for the state of Himachal Pradesh and would be located within the audio folder for Type A data.
● gbif_usage_key: this column contains a .json field with usage keys for mapping to the GBIF backbone taxonomy.
● media_info : a .json field containing media specific information such as sampling rate, bit depth.

For Type B data, the metadata.csv included the aforementioned columns for Type A and the following columns and associated naming conventions:

● scientific_name: contained both genus and species name, separated by an underscore, or only the highest rank when the species name was not provided.
● validation: this column contains 1, 0, or NA, indicating whether the recording passed validation, failed validation, or was not selected for validation.
● validation_fail_reason: this column contained specific reasons for why the recording failed validation. Poor audio quality, and misclassification of the recording (e.g. species not correctly identified) were the most common reasons identified for Type B data.
● validation_comments: specific notes provided by the validator to indicate why the recording was successful or unsuccessful at the validation stage.

For Type A data alone, an annotations.csv file was created and the following data naming conventions were used for the columns present:

● media_file_name: the unique file name corresponding to the audio file and contains the exact same information as mentioned in the corresponding column within the metadata.csv file.
● begin_time: the start time of the annotation in seconds.
● end_time: the end time of the annotation in seconds.
● high_freq: the highest frequency of the annotation in kilohertz.
● low_freq: the lowest frequency of the annotation in kilohertz.
● scientific_name: contained both genus and species name, separated by an underscore, or only the highest rank when the species name was not provided.
● validation: this column contains 1, 0, or NA, indicating whether the selection box drawn around the annotations passed validation, failed validation, or was not selected for validation.
● validation_fail_reason: this column contained specific reasons for why the annotation failed validation. Poor audio quality, misclassification of the recording (e.g. species not correctly identified), and annotation bounding box errors were the most common reasons identified for Type A data.
● validation_comments: specific notes provided by the validator to indicate why the annotation was successful or unsuccessful at the validation stage.
● notes: supplementary information provided by the authors. This field typically contained information on the call type, behaviour, and other observations of the organism.

The background data folders contained the same structure as above and included all sounds that did not contain the species of interest. The following columns, however, were not specified in either the metadata.csv or the annotations.csv associated with the background data folders: scientific_name, validation, validation_fail_reason, gbif_usage_key, and validation_comments. Instead, these columns were replaced with a single background_data column that indicates whether the recording or annotation contains wind, traffic, human vocalizations, or other background noise.

### Data Overview

Following data processing and technical validation (see section below), we found that the highest number of minutes of Type A (996 minutes) and Type B (1328 minutes) data came from Kerala and Madhya Pradesh, respectively (Figure 2). The southern Western Ghats montane rain forests recorded the highest proportion of Type A data (42.73 %; 1415 minutes), while the East Deccan moist deciduous forests of central India recorded the highest proportion of Type B data (53.30%; 1334 minutes) (Figure 3). In Type A data, 35,426 (96.48 %) out of the 36,718 annotations that contained organism names are at the level of species (including subspecies and domestic forms). In Type B data, a single audio file may contain multiple organisms (identified at the species level or highest possible rank). As a result, we created multiple rows (records) for the same audio file, each representing a unique scientific name. 27,297 (99.42 %) out of the 27,455 records (rows) were identified at the level of species (including subspecies and domestic forms) (Figure 4, 5).

**Figure 3.**
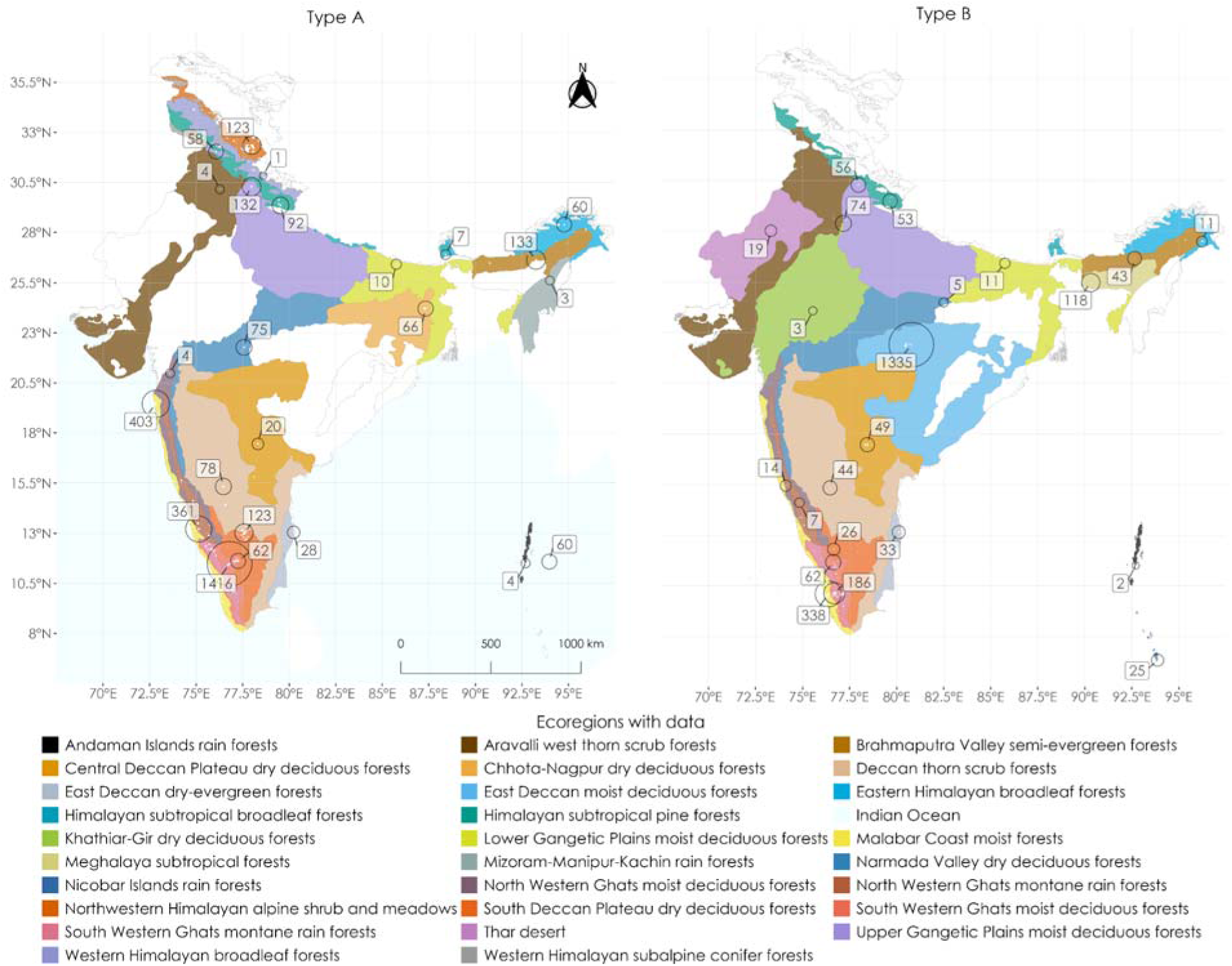
Bubble plot maps showing the duration of recordings (in minutes) for ecoregions in India (as well as the Indian Ocean), for both Type A and Type B data. The size of the bubble indicates the total duration of recordings submitted from the corresponding ecoregion. Two Type B recordings (around 71 seconds in total media length) with no geographic information are not included in the display. White spots represent the coordinates provided for each recording (they may not be exact). For each ecoregion, bubbles are centered around locations with the maximum number of recordings (indicated by the black arrow), except the Indian Ocean for the purpose of visibility. Map not for scale.

**Figure 4.**
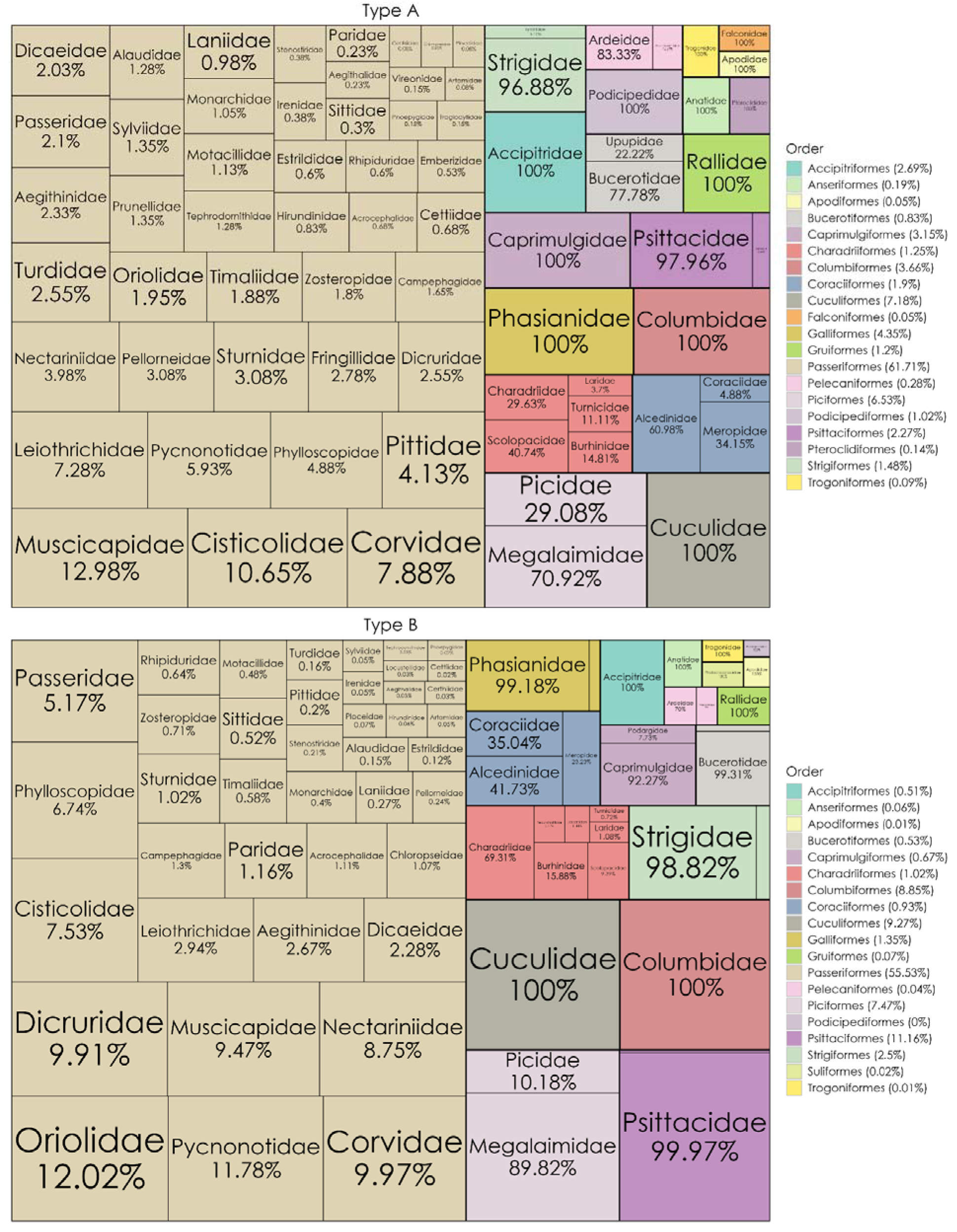
Tree map showing the proportio ns of label s of avian taxa alone (at the order and family level). The percentage s represent the num ber of label s of birds falling within a given family divided by the number of labels of birds falling in that order under which the family belongs. The size of the rectangles represents the proportion of labels that belong to a particular order. The combined size of the same-coloured rectangles represent the proportion of labels falling within that specific order (this percentage proportion, the name of the order and the corresponding colour are displayed in the legend on the right). To make the data comparable between both types, even if multiple annotation labels exist for a given species within the same recording, only one of those labels for that species is retained for Type A (it is by default the case in Type B).

**Figure 5.**
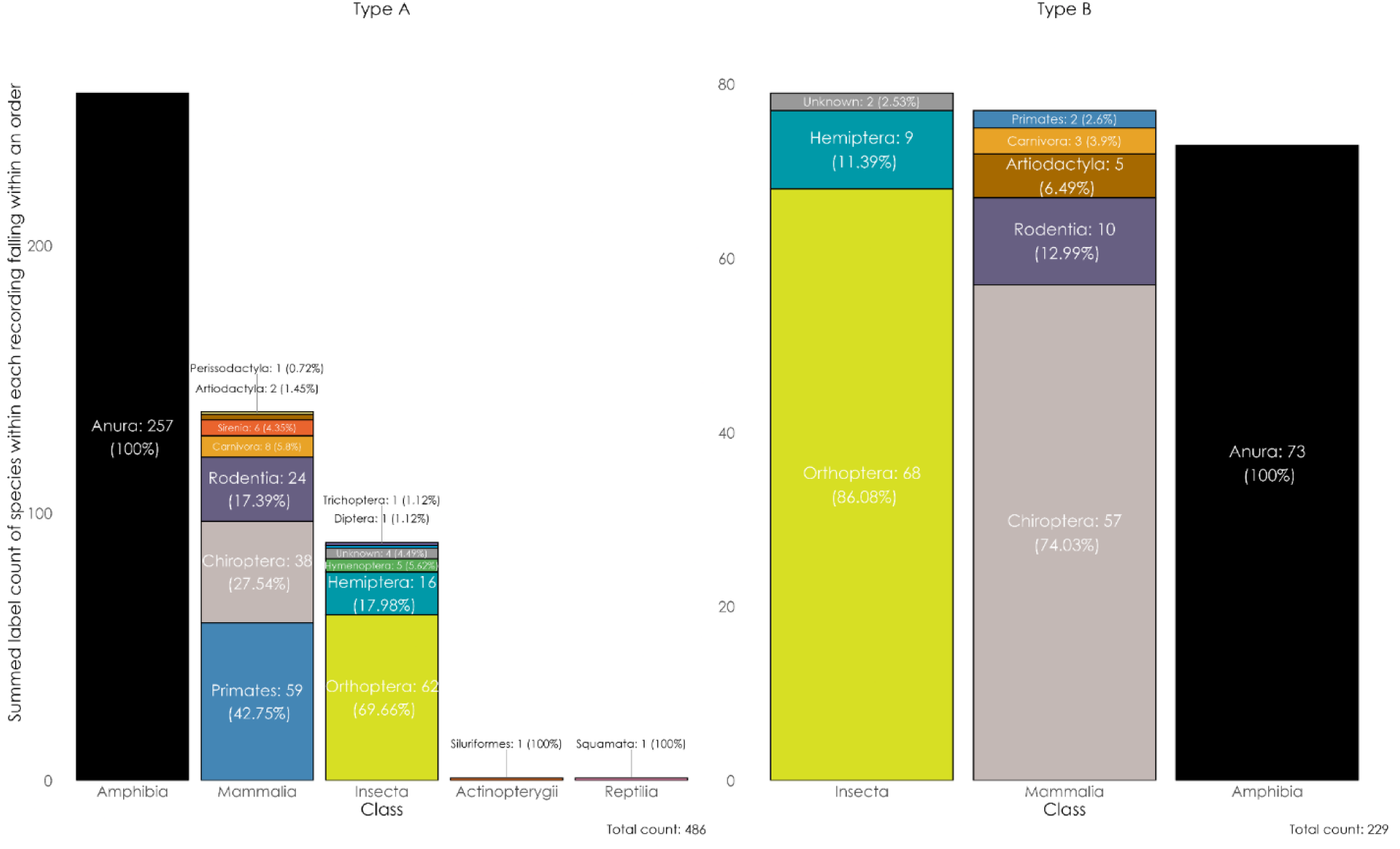
This stacked bar-plot shows the taxonomic (class- and order-wise) breakdown of the non-avian taxa in both Type A and Type B data. The numbers alongside the order names represent the number of organism labels which fall within that order across all audio files. To make the data comparable between both types, even if multiple annotation labels exist for a given species within the same recording in Type A data, only one of the labels for that species is retained.

### Technical Validation

For technical validation, we first created a validation dataset by randomly sampling 2% of the total data for each state-taxonomic class combination, separately for Type A and Type B data. Since the submitted data disproportionately represented more bird species and certain geographical regions (such as the Western Ghats and Western Himalayas), we chose the above sampling approach to ensure fair weightage across classes and regions during validation. For Type A data, we requested two expert validators to verify all annotations in each sampled file and for Type B data, we requested the two expert validators to verify all records associated with a sampled file.

We first anonymised the file names of the chosen audio files, and only the location, recording date, and notes column were shown to the validators to reduce any biases. We uploaded these files into a folder for two independent validators to access. Neither validator was involved in the creation, submission, or processing of any of the datasets. Table 1 and 2 shows the total number of media files sampled for validation for Type A and Type B data by each state and taxonomic class combination, and associated percentages of data that did not pass validation.

**Table 1.** Type A validation data and results, with the total number of annotations (labels) and files that failed validation, and the associated reasons.

| State | Taxonomic Group | #Media Files | #Labels | #Fail Labels | #Fail Media Files | % Fail Media Files | % Fail Labels |
| --- | --- | --- | --- | --- | --- | --- | --- |
| Andaman and Nicobar Islands | Aves | 1 | 15 | 0 | 0 | 0.0% | 0.0% |
| Andhra Pradesh | Aves | 2 | 10 | 1 | 1 | 50.0% | 10.0% |
| Andhra Pradesh | Mammalia | 1 | 3 | 0 | 0 | 0.0% | 0.0% |
| Arunachal Pradesh | Aves | 1 | 5 | 0 | 0 | 0.0% | 0.0% |
| Assam | Aves | 1 | 5 | 0 | 0 | 0.0% | 0.0% |
| Bihar | Aves | 2 | 16 | 5 | 1 | 50.0% | 31.3% |
| Bihar | Insecta | 1 | 3 | 3 | 1 | 100.0% | 100.0% |
| Bihar | Mammalia | 1 | 18 | 0 | 0 | 0.0% | 0.0% |
| Goa | Aves | 1 | 16 | 16 | 1 | 100.0% | 100.0% |
| Gujarat | Aves | 1 | 20 | 2 | 1 | 100.0% | 10.0% |
| Haryana | Aves | 1 | 5 | 0 | 0 | 0.0% | 0.0% |
| Himachal Pradesh | Amphibia | 1 | 13 | 0 | 0 | 0.0% | 0.0% |
| Himachal Pradesh | Aves | 6 | 126 | 0 | 0 | 0.0% | 0.0% |
| Himachal Pradesh | Insecta | 1 | 1 | 0 | 0 | 0.0% | 0.0% |
| Himachal Pradesh | Mammalia | 1 | 15 | 0 | 0 | 0.0% | 0.0% |
| Jammu and Kashmir | Aves | 1 | 32 | 0 | 0 | 0.0% | 0.0% |
| Jharkhand | Aves | 2 | 31 | 1 | 1 | 50.0% | 3.2% |
| Karnataka | Amphibia | 1 | 2227 | 0 | 0 | 0.0% | 0.0% |
| Karnataka | Aves | 5 | 122 | 28 | 3 | 60.0% | 23.0% |
| Karnataka | Insecta | 1 | 15 | 0 | 0 | 0.0% | 0.0% |
| Karnataka | Mammalia | 2 | 23 | 0 | 0 | 0.0% | 0.0% |
| Kerala | Amphibia | 1 | 12 | 0 | 0 | 0.0% | 0.0% |
| Kerala | Aves | 2 | 115 | 0 | 0 | 0.0% | 0.0% |
| Kerala | Insecta | 2 | 63 | 0 | 0 | 0.0% | 0.0% |
| Kerala | Mammalia | 1 | 4 | 0 | 0 | 0.0% | 0.0% |
| Kerala | Squamata | 1 | 1 | 0 | 0 | 0.0% | 0.0% |
| Maharashtra | Aves | 7 | 393 | 0 | 0 | 0.0% | 0.0% |
| NCT of Delhi | Mammalia | 1 | 5 | 0 | 0 | 0.0% | 0.0% |
| Nagaland | Aves | 1 | 67 | 0 | 0 | 0.0% | 0.0% |
| Punjab | Aves | 1 | 4 | 0 | 0 | 0.0% | 0.0% |
| Tamil Nadu | Amphibia | 1 | 21 | 0 | 0 | 0.0% | 0.0% |
| Tamil Nadu | Aves | 1 | 55 | 0 | 0 | 0.0% | 0.0% |
| Tamil Nadu | Mammalia | 2 | 2 | 0 | 0 | 0.0% | 0.0% |
| Uttarakhand | Aves | 1 | 14 | 0 | 0 | 0.0% | 0.0% |
| Uttarakhand | Mammalia | 1 | 9 | 0 | 0 | 0.0% | 0.0% |
| **Total** |  | **57** | **3486** | **56** | **9** | **16%** | **2%** |

| **Type A Validation Fail Reasons** |  |  |
| --- | --- | --- |
| **Reason for Validation Fail** | **#Media Files** | **#Labels** |
| Audio-related | 2 | 3 |
| Bounding-Box-related | 1 | 2 |
| Label-related | 8 | 53 |
| Reviewer-related | 0 | 0 |
| Taxa-related | 0 | 0 |

**Table 2.**
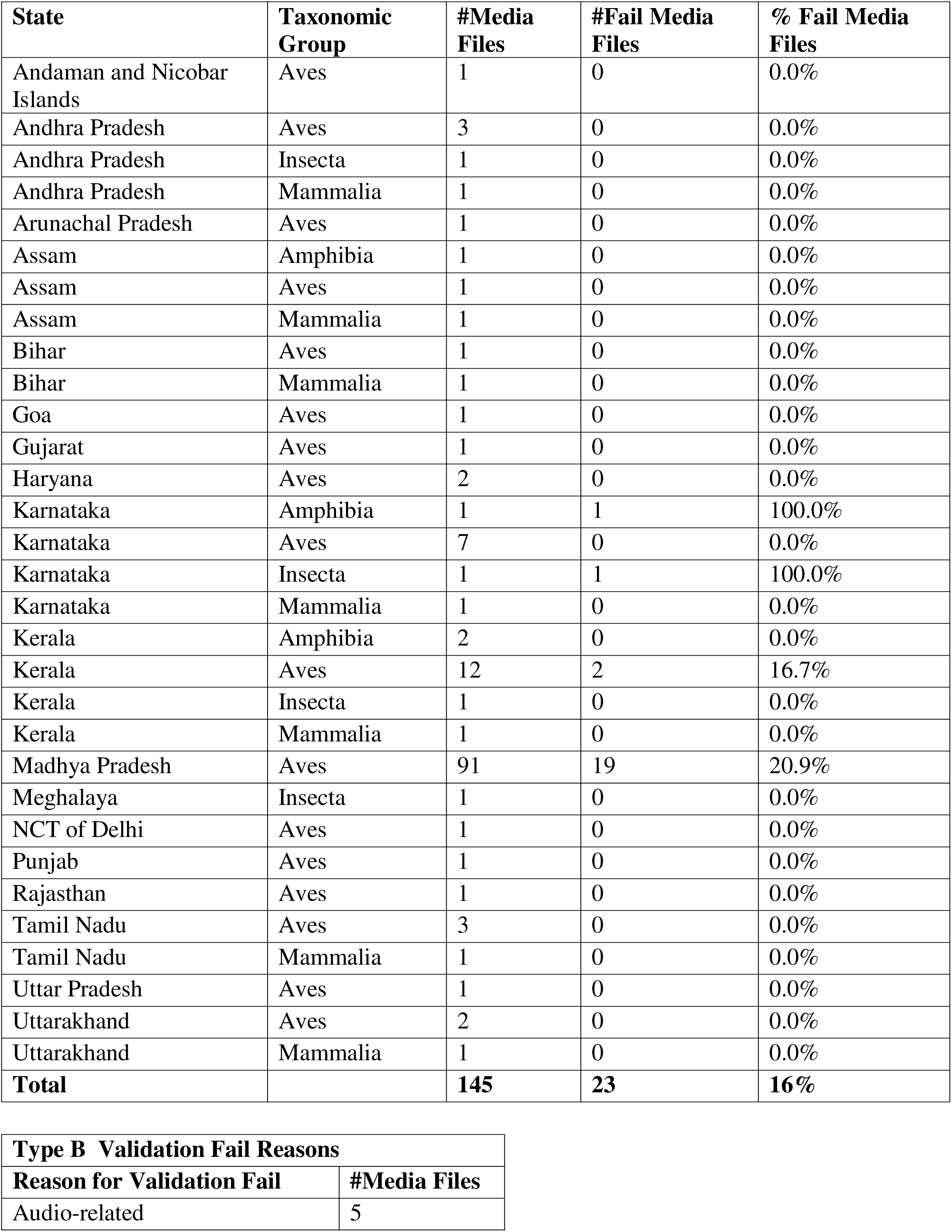
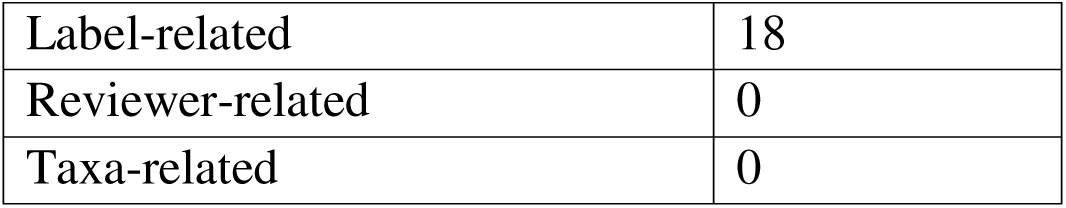
Type B validation data and results, with the total number of files that failed validation, and the associated reasons.

To ease the process of validation, we relied on a lightweight JavaScript application called Annotation Viewer^18^, which could be used by all reviewers to visualise the spectrogram of a selected audio file along with annotations (if present). The validator would then tick the Pass/Fail checkbox and insert their comments on the application (Figure 6). This software also ensured that the validation process was consistent across the two validators.

**Figure 6.**
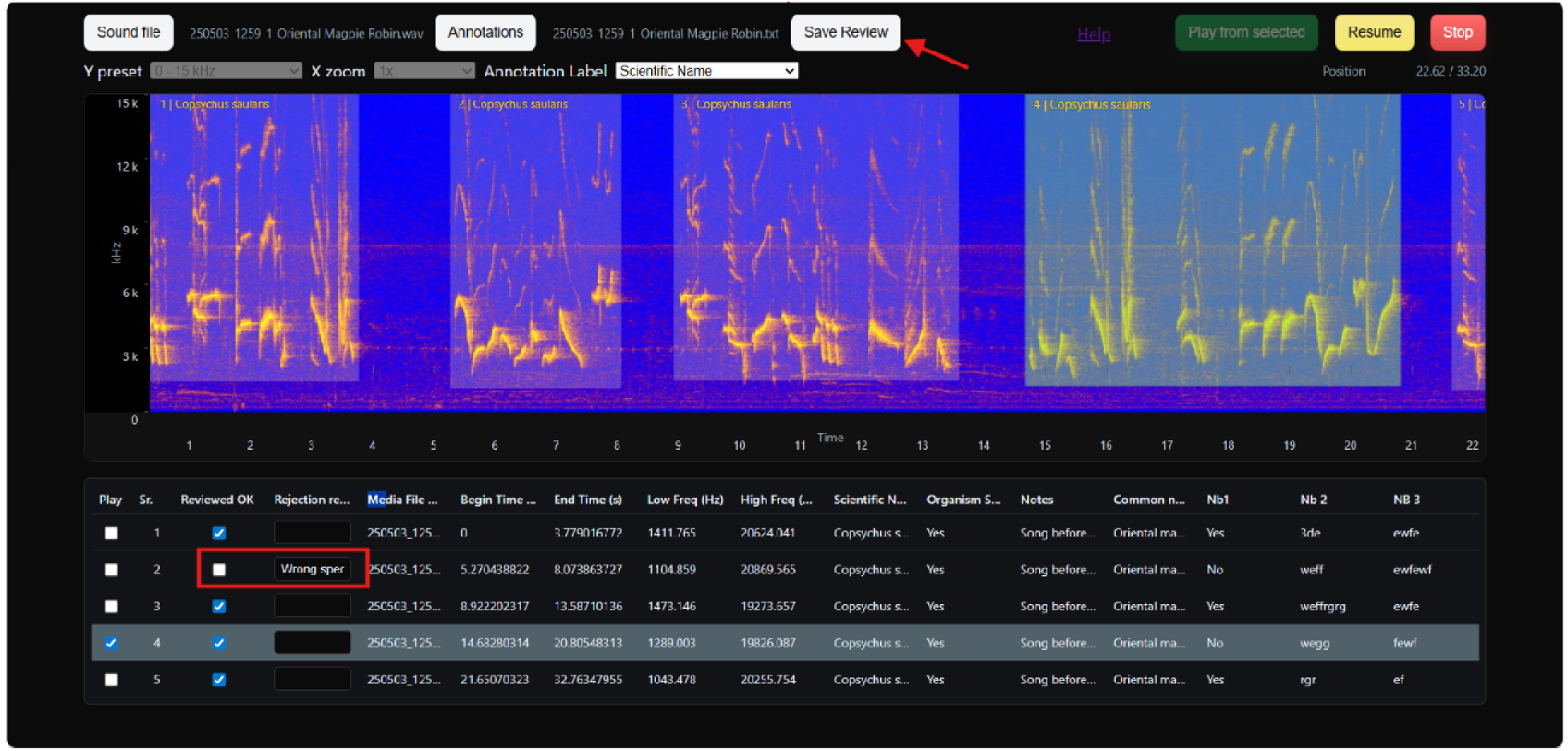
Annotation Viewer was used for technical validation. The validator marked each annotation (if present) in a recording as OK or rejected them with associated comments.

Following the process of technical validation, we obtained an .xlsx file containing the media file names, a validation column with a 1 or 0 indicated by the validator to suggest if the audio file passed or failed validation, a validation_fail_reason column containing broad categories for why an audio recording failed validation (categories include label-related and audio-related), and a validation_comments column which contained any detailed information provided by the validators for why they assessed a particular recording as unsuccessful or successful. Initially, validators also marked files as having failed validation for two reasons (as noted in the validation_comments column): a) expertise related: files that had taxonomic groups that required high levels of expertise (e.g., insects, bats, freshwater mammals) and such species had limited reference acoustic templates for confirming their identity and b) data processing related: files that contained the appropriate scientific name but were mapped under the wrong class while programmatically aligning with the GBIF backbone.

Since expertise related failures were associated with files contributed by taxonomic experts, we chose to mark these files as having passed validation and such changes have been indicated in the validation_comments column in the Zenodo dataset. In the case of data processing failures, we reviewed the entire dataset, including the validation files, corrected spelling errors, and ensured that the correct class was assigned to each scientific record.

Across Type A data, a total of 3486 annotations from 57 files were sampled for validation, of which 98% of annotations passed validation while 9 files had at least one annotation that failed validation. Of the 56 annotations that failed, 53 failed due to labeling issues (species were misidentified), 3 failed due to audio quality issues (faint or poor audio), and 2 failed due to errors in the placement of the bounding boxes around the annotation. Across Type B data, a total of 145 files were sampled for validation, of which 84% passed validation. 18 failed due to labeling issues (species misidentification) and five files were of poor audio quality in which a validator could not identify the species within that file.

## Usage Notes

### Dataset challenges

One of the biggest challenges and strengths of this dataset is that our annotated data came from a variety of sources contributed by participatory scientists, researchers, conservation practitioners, and nature enthusiasts. Given the variation in how the data were annotated and collected, we faced several challenges in processing this dataset. First and foremost, expecting all participants to provide the same audio format and follow the exact same protocol for annotation was found to be impractical. As a result, we spent time processing and organizing the data so that it is usable and readable when accessed via Zenodo. Secondly, despite our efforts to validate data, we acknowledge that minor annotation errors may persist across datasets and taxonomic groups.

However, we are confident that these errors are minimal and contain valuable information for future machine learning research and development of automated recognition algorithms. Lastly, we acknowledge that our dataset is a biased representation of taxonomic diversity in India, as the majority of contributed annotations were for avifauna. However, our hope is that future iterations of the dataset creation will place a targeted emphasis on taxonomic groups such as insects, bats, and marine and freshwater fauna. Our data descriptor and the associated dataset showcase the strengths of participatory science, and we believe that to solve urgent problems in conservation biology, we require novel, unique approaches, such as ours.

## Data and code availability

The acoustic data and associated annotations (Type A, Type B and background data) are hosted in Zenodo under the CC-BY-NC license. For Type A data, please visit: https://zenodo.org/records/18743214. For Type B data, please visit: https://zenodo.org/records/18927866. For background data, please visit: https://zenodo.org/records/18928201. All the associated code for data pre-processing, validation, and figure generation are available in these GitHub repositories: https://github.com/paulvpop/supporting-code-for-A-large-scale-crowd-sourced-acoustic-dataset-of-Indian-fauna and https://github.com/chimarkhi/ien-data-mgmt<u>#</u> under the MIT license. The google form associated with uploading data and other ancillary material can be accessed here: https://forms.gle/PBybpPnrRR6YZJEU9

## Author contributions

V.R., S.S., P.P., P.C., P.S., S.K., S.T., P.B., and K.D. were involved in conceptualization, methodology iteration, and scientific communication. S.S. and P.P. led the analysis. V.R. acquired funding. V.R. and S.S. coordinated the project. V.R., S.S., P.P., P.S., S.K., and S.T. were involved in capacity building (annotation workshops and office hours). S.S., P.P., and P.C., were involved in the design of the validation process. P.P., P.S., and S.K. were involved in creating visualizations. V.R. led the writing. All co-authors contributed to the submission of data and reviewed the writing of the manuscript.

## Acknowledgements

We would like to thank all members of the India Ecoacoustics Network for their contributions of data and time to this effort. We also acknowledge Vijay Kumar, Krishna Kumar, Nature Conservation Foundation, Divya Mudappa, T R Shankar Raman, Anagha C S, Jishnu Borgohain, Anand M. Osuri, Rohit Naniwadekar, Vignesh Poojary, Kiran, and Vidwath S M for their assistance with data collection. We thank Devendra Korche for assistance with data collection in Mandla, Madhya Pradesh.

